# Aperiodic exponent ceases to track changes in propofol dose at deep surgery-level hypnotic depths

**DOI:** 10.64898/2026.06.17.732951

**Authors:** Gonzalo Boncompte, Juan Cristóbal Pedemonte, Luis I. Cortínez, Mauricio Ibacache, Natalia Caleron, Daniela Biggs

**Affiliations:** División de Anestesiología, Escuela de Medicina, Pontificia Universidad Católica de Chile; Red de Salud UC CHRISTUS, Santiago, Chile; Programa de Farmacología y Toxicología, Facultad de Medicina, Pontificia Universidad Católica de Chile

**Keywords:** EEG, Anaesthesia, aperiodics, scale-free EEG, excitation/inhibition balance, cortical states

## Abstract

**Background:** Accurately tracking hypnotic depth during surgery is key to deliver personalized and precise anaesthesia. The aperiodic exponent of electroencephalographic (EEG) data has shown great promise as a marker of the balance between excitatory and inhibitory cortical activity and of hypnotic depth. In humans, this relation has been mostly evidenced by pairwise comparisons between awake and superficial anaesthesia states. However, it is unclear if this relation is also present across deeper hypnotic depths. Here we examined the relation between aperiodic exponent and hypnotic depth across different surgery-level propofol doses.

**Methods:** The present was an exploratory prospective observational analytical study where we recorded high-density EEG data from patients before surgery and during continuous target-controlled propofol infusions with different intraoperative target concentrations. We evaluated the relation between steady-state propofol dose and several EEG features including aperiodic exponent and spectral band powers.

**Results:** All EEG measures were different when comparing awake versus anaesthesia conditions. At surgery-level hypnotic depths, some spectral band powers continued to track propofol dose displaying either with positive (delta R^2^ = 0.23, *p* = 0.001) or negative correlations (alpha R^2^ = 0.36, p < 0.001; beta R^2^ = 0.045, p < 0.001). Importantly, the aperiodic exponent was not associated with propofol dose’s absolute value (R^2^ = 0.011, *p* = 0.43) or with its relative change (R^2^ = 0.011, *p* = 0.51).

**Conclusions:** Our results suggest that the relation between aperiodic exponent and excitatory/inhibitory balance is circumscribed to more superficial hypnotic depths than those typically employed for surgery.

## Introduction

A key goal of precision medicine in the context of anaesthesia is that doses should be chosen to produce a specific effect in each patient, rather than with a one-fits-all approach.^1^ This is of particular relevance given the known negative effects of both excessive ^2^ and insufficient ^3^ doses of hypnotic drugs during surgery. To achieve this, monitoring of brain activity intraoperatively with electroencephalography (EEG) can help directly evaluate the effects of hypnotic drug on their target organ. However, the reliability and effectiveness of EEG monitoring ultimately depend on our ability to understand and precisely model the relation between brain activity and hypnotic depth.

For nearly 100 years now, the effect of inhibitory hypnotic drugs on cortical electrical activity has been known: the classic “slowing down” of the EEG signal, meaning a reduction in the frequency of the most prominent oscillations present.^4^ More recent evidence has supported and given further details about this, showing that delta/slow frequency bands systematically increase in response to, for example, GABAergic drugs, while high frequency bands (e.g. beta and gamma ranges) are reduced.^5, 6^ These known relations are leveraged by current commercial EEG monitors to deliver numerical estimates of hypnotic depth.^7, 8^ However, current EEG monitors have known biases and shortcomings, especially in vulnerable populations.^9, 10^ Part of the reason for this is the relatively poor neurobiological/theoretical background that supports them, which makes research on this topic particularly relevant.

Recent developments in theoretical and computational neuroscience have advanced the concept of the aperiodic component as a valuable tool to describe EEG signals and cortical states.^11–15^ In this framework, the frequency spectrum is evaluated as a whole instead of frequency band by frequency band, by adjusting a power law function to its power spectral density and evaluating the resulting aperiodic exponent. ^16^ Computational models have suggested that this parameter monotonically tracks the balance between excitatory (glutamatergic) and inhibitory (GABAergic) activity (E/I Balance). This has been supported by evidence from rats, monkeys, and humans under anaesthetics where baseline and anaesthetised conditions are compared. They systematically show that GABAergic drugs produce an increase in the aperiodic exponent ^14, 17^ (steeper slope). In fact, Widmann and cols (2025) ^18^ recently showed that the aperiodic exponent has a greater discriminatory power than several spectral measures to differentiate awake and anaesthetized states in human patients.

Importantly, most of these studies compare awake and anaesthesia states without consideration for the intermediate states between fully awake and deep surgery-level anaesthesia. Understanding the full trajectory of these measures is key if they are ever to be included in clinically relevant EEG monitors. This is especially important because we know that several EEG features can show non-linear non-monotonic relations to hypnotic depth including alpha power ^5^ and Lempel-Ziv complexity ^19^. In the present study we directly evaluated how well the aperiodic exponent tracks different surgery-level propofol doses, as well as in comparison to baseline (awake) levels. We compare its performance to that of canonical spectral bands power.

## Methods

The present was an exploratory prospective observational analytical study. It was conducted in the Hospital Clinico UC-Christus with the approval of the local ethics committee (CEC Salud UC N° 190912008) and in accordance with the guidelines of the Helsinski declaration.

### Patients

We recruited a total of 17 patients from the Hospital Clinico UC-Christus from 2022 to 2024. Patients scheduled for surgery were informed of the study protocol and invited to participate. Inclusion criteria comprehended patients between 18 and 55 years old, ASA Physical status I or II, scheduled for a non-neurological surgery with an estimated duration between 1.5 and 3 hours and with an anaesthetics plan of propofol-remifentanil total intravenous anaesthesia. These drugs were delivered using targeted-controlled infusions (TCI) with the Marsh and Minto models, respectively. Exclusion criteria were epilepsy, major depression, schizophrenia, or a history of stroke. The scheduled use of dexmedetomidine was also an exclusion criterion.

### EEG recordings

EEG was recorded using a high-density 64-channels Neuroscan system both before and during surgery with 2kHz sampling frequency with positioning according to the 10-20 international system. Baseline EEG recordings were conducted with closed eyes for 5 minutes in a sitting resting state before any anaesthetics were delivered. Intraoperative EEG was recorded from the time of induction until the recovery of verbal responses at the end of the surgery.

### Intraoperative steady-state periods

We aimed to evaluate different EEG markers of hypnotic depth at stable Marsh’s effect-site propofol concentrations (Ce). For this, we separated the surgery into discrete periods at which propofol Ce (Marsh model) was maintained at a constant value for at least 20 minutes. Based on previous literature,^20^ this would allow us to reach steady-state propofol Ce in 10 minutes of TCI, leaving at least 10 minutes of EEG recordings at each stable propofol Ce. During these steady-state periods, propofol concentration was (attempted to be) maintained at a particular value defined by clinical requirements according to the attending anaesthesiologist. If clinical conditions required dose adjustments, these changes were conducted and that steady-state’s recording was discarded. All other drugs were freely modified by the attending anesthesiologist at any moment according to clinical requirements. After each steady-state period (20-30 minutes) propofol Ce was modified for another steady-state period. This continued until the end of surgery or until the expected surgery duration did not allow for another 20 minutes of recording.

### EEG analyses

#### Preprocessing

All EEG recordings were analysed using custom-made Python scripts based on previous works and employing open-access libraries including numpy, scipy, matplotlib, specparam,^16^ and MNE.^21^ EEG data from intraoperative steady-state periods were extracted. All EEG data to be analysed were first visually inspected to discard electrodes and time periods with artifacts, for example when an electrosurgical unit was employed or for electrodes with poor electrical continuity.

#### Spectral band power estimations

Clean EEG data was segmented into 2.5s epochs (40% overlapping) using the multitaper strategy as implemented by the mne.time_frequency.psd_array_multitaper MNE function with full normalization. After calculating the power spectral density (PSD), the median power within each frequency band was calculated for the delta band (0.5 - 3.5Hz), theta band (4 - 7Hz), alpha band (7.5 - 12Hz) and beta band (13 - 25Hz). We also estimated the total power as the sum of the power of all frequency components between 0.5 and 40 Hz. Afterwards the spectral band powers were transformed from uV^2^/Hz into decibels and averaged across a broad frontal region of interest (electrodes FP1, FPZ, FP2, AF3, AF4, F11, F7, F5, F3, F1, FZ, F2, F4, F6, F8, F12, FT11, FC5, FC3, FC1, FCZ, FC2, FC4, FC6 and FT12) and across time for each steady-state period. The data from the first 10 minutes of each steady-state period were discarded.

#### Aperiodic exponent estimations

Aperiodic estimations were conducted using the same spectral estimations (multitaper, 2.5s window) and the specparam library (formerly known as FOOOF ^16^). We used a fixed aperiodic model (no knee), peak width limits between 1 and 5 Hz, 2 max peaks and a minimum peak height of 0.1 dB. Models with a poor fit (R^2^ < 0.8) were discarded from further analyses. After aperiodic model estimation was conducted separately on each time point and electrode the aperiodic exponent, offset and R^2^ (goodness of fit) values were averaged across time and electrodes for each steady-state period.

### Statistical analyses

Differences between EEG measures in baseline and anaesthesia were evaluated using paired t-tests. The dependence between target propofol concentration and intraoperative EEG measures (band powers and aperiodic exponent) were assessed by means of Pearson’s correlations. Outliers in the differences between the spectral band power or aperiodic exponent of consecutive steady-state periods were classified as such based on an absolute threshold of 3 (z-score) and discarded from further analyses.

## Results

We recorded EEG activity from a total of 17 patients before anaesthesia (baseline) during a closed-eyes awake resting state and during intraoperative periods with stable target effect-site propofol concentrations (propofol Ce). Each patient went through between 2 and 6 intraoperative periods lasting at least 20 minutes each, producing a total of 60 steady-state periods. We discarded the first 10 minutes of each period to ensure a steady state propofol Ce during the whole analysed time. Stable intraoperative propofol Ce varied between states between 2.3 µg*ml^-1^ and 5.0 µg*ml^-1^, with an average of 3.27 µg*ml^-1^ (s.d. = 0.6). Patient’s age ranged between 21 and 57 years (average = 41.8, s.d. = 12.6). EEG data were visually inspected for artifact rejection, eliminating on average 17.5% (s.d. 12.2%) of time per period.

### Spectral EEG measures in baseline vs anaesthesia

We estimated EEG features in baseline and intraoperative time periods, including spectral band powers and the aperiodic exponent (see Figure 1 for representative PSDs and aperiodic fits). We compared the median spectral power within canonical frequency bands during baseline and during each patient’s first intraoperative period. In this binary comparison, propofol anaesthesia significantly increased the spectral power within all frequency bands (Figure 2; delta t(16) = 10.5, p<0.001; theta t(16) = 6.5, p<0.001; alpha t(16) = 5.4, p<0.001; beta t(16) = 2.6, p = 0.020; total power t(16) = 10.3, p<0.001). Aperiodic exponent also increased (steeper slope) from baseline to anaesthesia (aperiodic exponent t(16) = 15.0, p < 0.001; Figure 2). The average Ce for patient’s first intraoperative period was 3,26 µg*ml^-1^ (s.d. 0.72).

**Figure 1.**
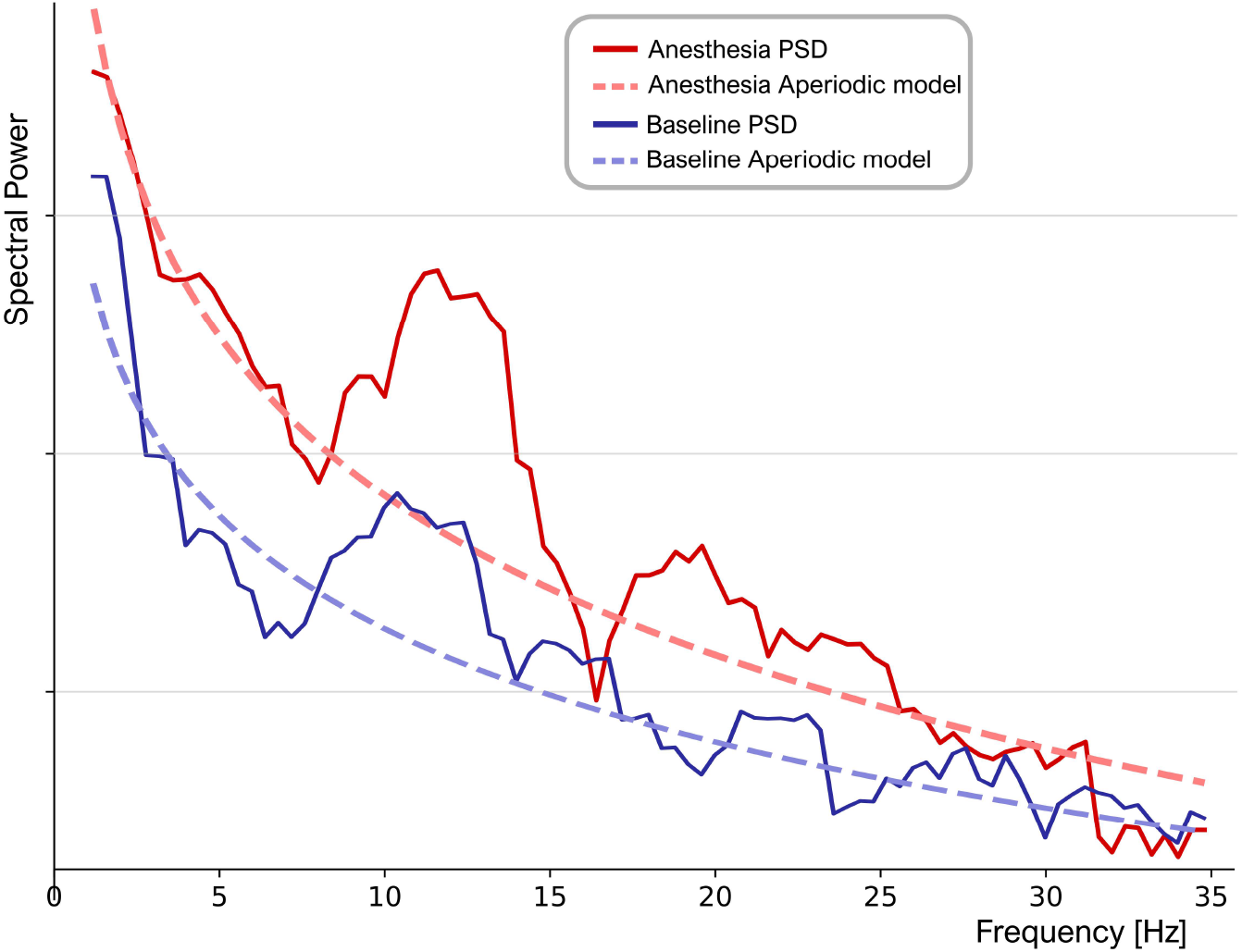
Representative PSDs of Baseline and anaesthesia periods with their respective best aperiodic power-law fit.

**Figure 2.**
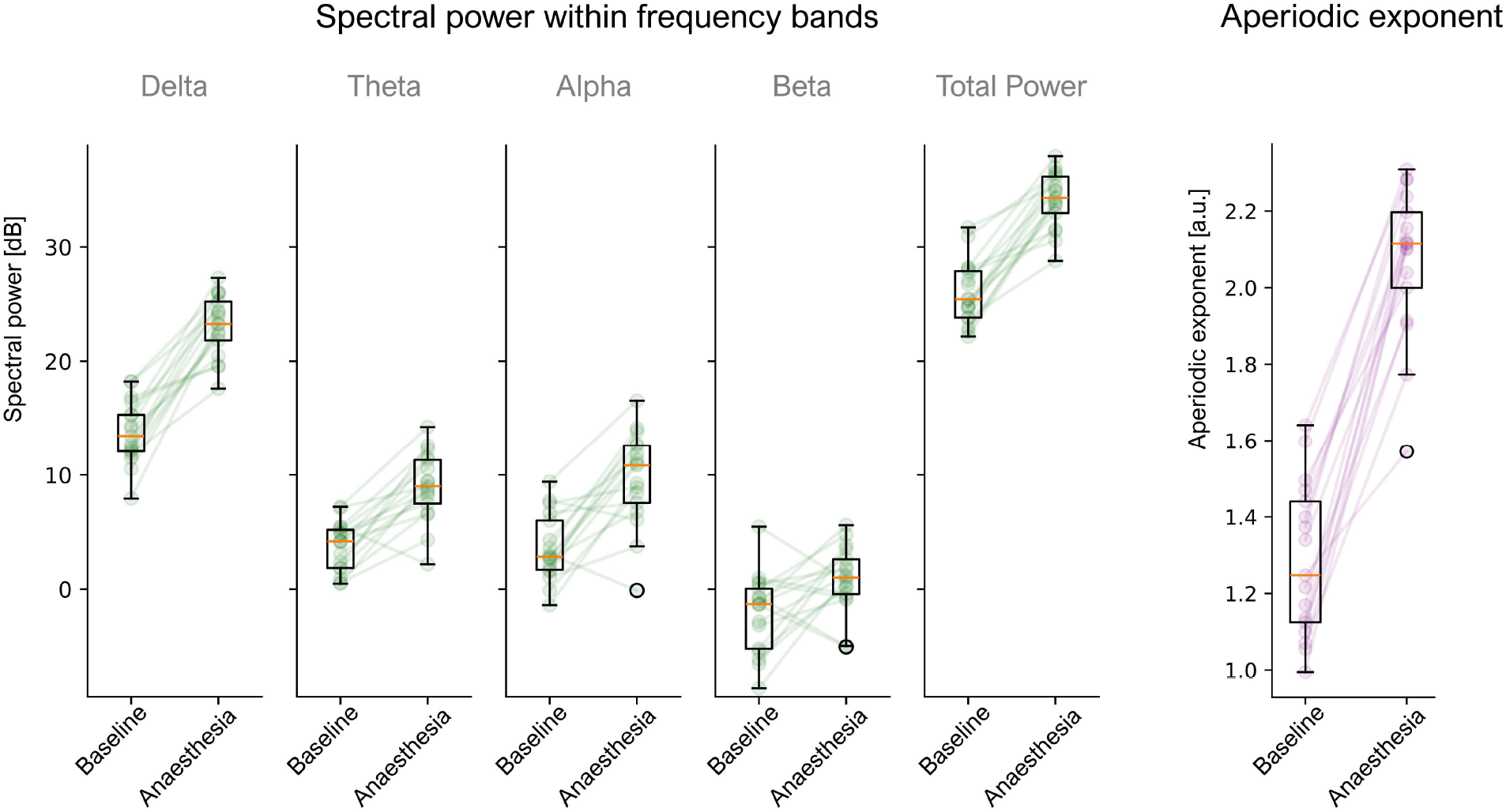
Changes in EEG measures between awake and anaesthesia. Boxplots depicting the comparisons between EEG measures during baseline and the first steady-state intraoperative period. EEG measures include spectral band powers (green) and aperiodic exponent (purple).

### Correlations between propofol dose and spectral measures in absolute values

We next evaluated continuously the relation between each EEG measure and propofol Ce across different intraoperative periods. We found significant positive correlations for the delta frequency band (Figure 3; r = 0.32, p = 0.014) and for the total band power (r = 0.28, p = 0.032). Theta, alpha and beta bands did not show a significant correlation with propofol Ce. Interestingly, the aperiodic exponent also did not show a significant correlation with propofol dose (r = 0.10, p = 0.43).

**Figure 3.**
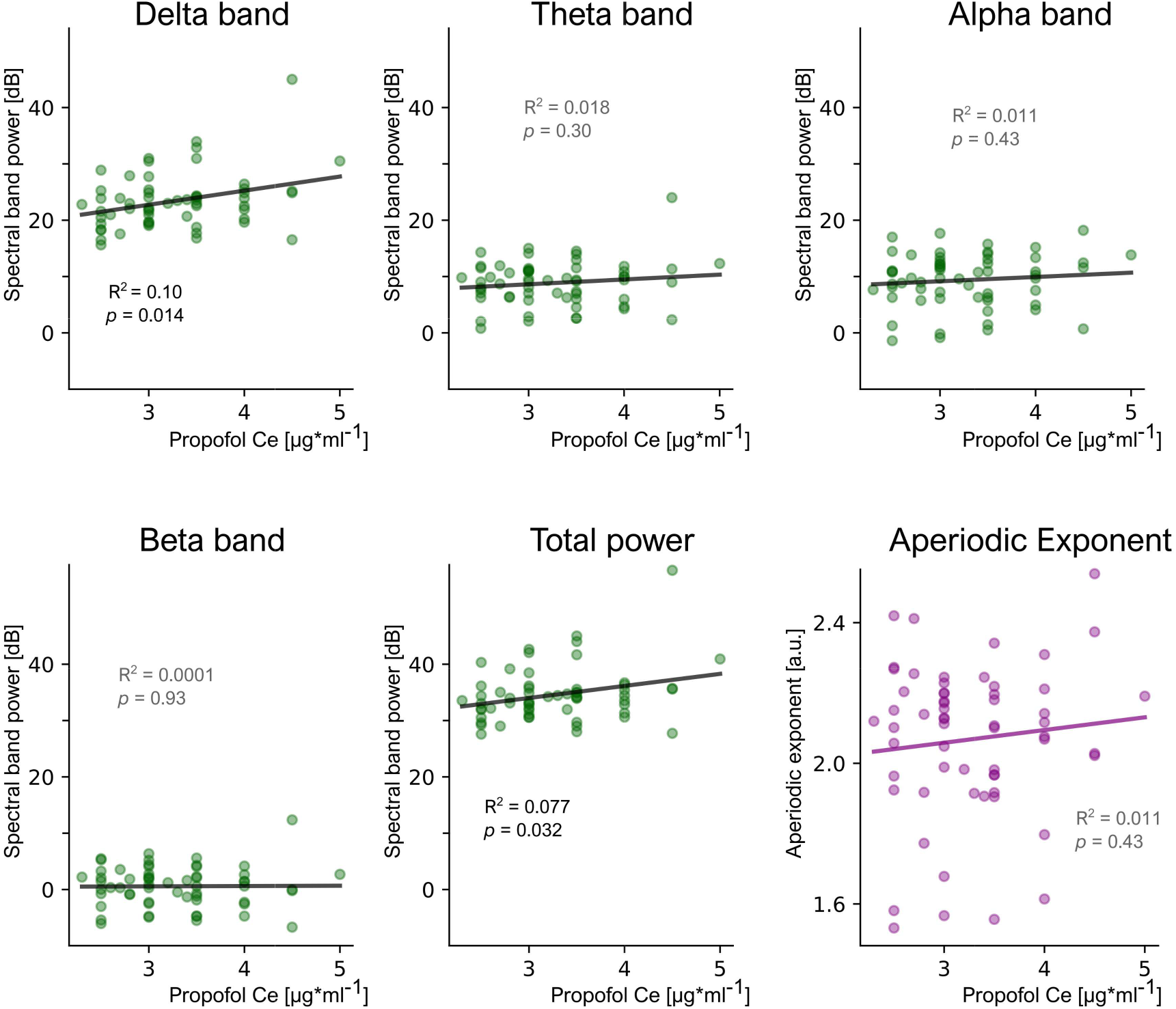
Relations between absolute of EEG measures and the absolute propofol effect site concentrations. Scatter plots showing the associations between EEG measures (spectral band powers in green, aperiodic exponent in purple) and target propofol concentrations for all steady-state intraoperative periods. Black lines show the best linear fit for each spectral measure. R^2^ and *p* values correspond to Pearson’s correlation tests. R^2^ and p values in grey indicate non-significant correlations.

### Associations between changes in propofol doses and changes in EEG measures in transitions between intraoperative periods

Because the same propofol Ce can have markedly different effects on patients depending on their individual characteristics, we evaluated the relative modulations of EEG measures in response to changes in propofol Ce. For each pair of consecutive intraoperative periods, we calculated the change in each EEG measure and in propofol Ce (Figure 4). Consistent with previous analyses, we found significant positive correlations for delta band (r = 0.48, p = 0.0015) and total power (r = 0.40, p = 0.009). Changes in propofol Ce were also negatively associated with spectral power in the alpha and beta bands (alpha r = -0.60, p < 0.001; beta r = -0.67, p < 0.001). Importantly, changes in target propofol concentration were not associated with relative changes in the aperiodic exponent of the EEG signal (r = 0.10, p = 0.51).

**Figure 4.**
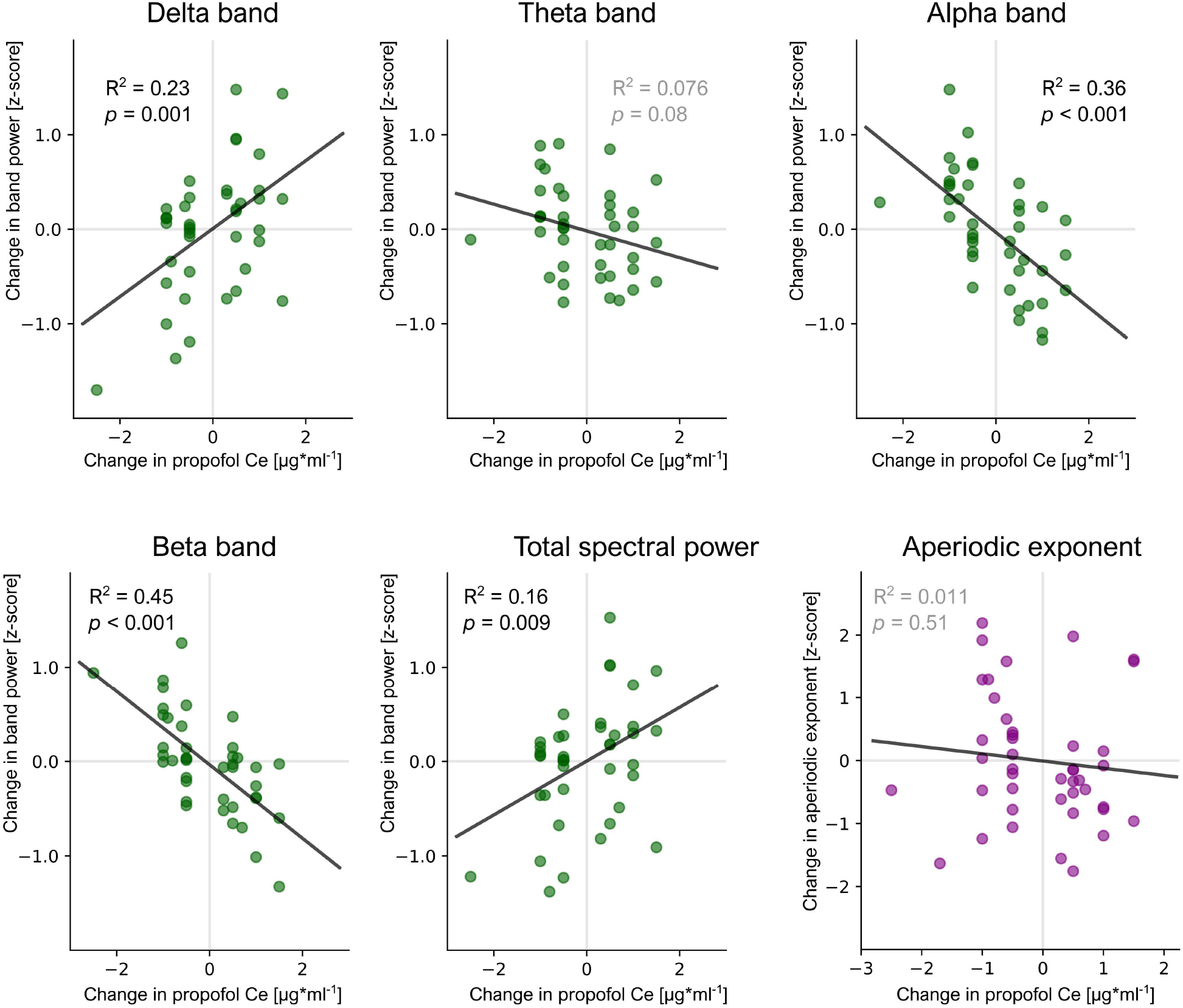
Relation between changes in periodic and aperiodic measures versus changes in propofol dose. Scatter plots showing the associations between the amount of change in spectral measures versus the amount of target propofol concentration change between consecutive steady state intraoperative periods. Green disks represent band powers and purple ones represent aperiodic exponent. Each disk represents the difference between two consecutive intraoperative periods. Black lines show the best linear fit for each spectral measure. R^2^ and *p* values correspond to Pearson’s correlation tests. R^2^ and p values in grey indicate non-significant correlations

## Discussion

In the present work we inquired about the relation between EEG spectral measures and propofol dose at different hypnotic depths. We compared EEG’s band powers and aperiodic exponent between awake and anesthetized and also between different surgery-level doses of propofol anaesthesia. We found that delta band power consistently increased from awake to anaesthesia states in general and from moderate to deeper anaesthesia states. In contrast, the aperiodic exponent showed marked differences between awake and anaesthesia, but it was no longer sensitive to differences of propofol doses at surgery-level hypnotic depths. This is summarized in Figure 5, which displays a model of the trajectories of delta band power, alpha band power and the aperiodic exponent though different hypnotic depths.

**Figure 5.**
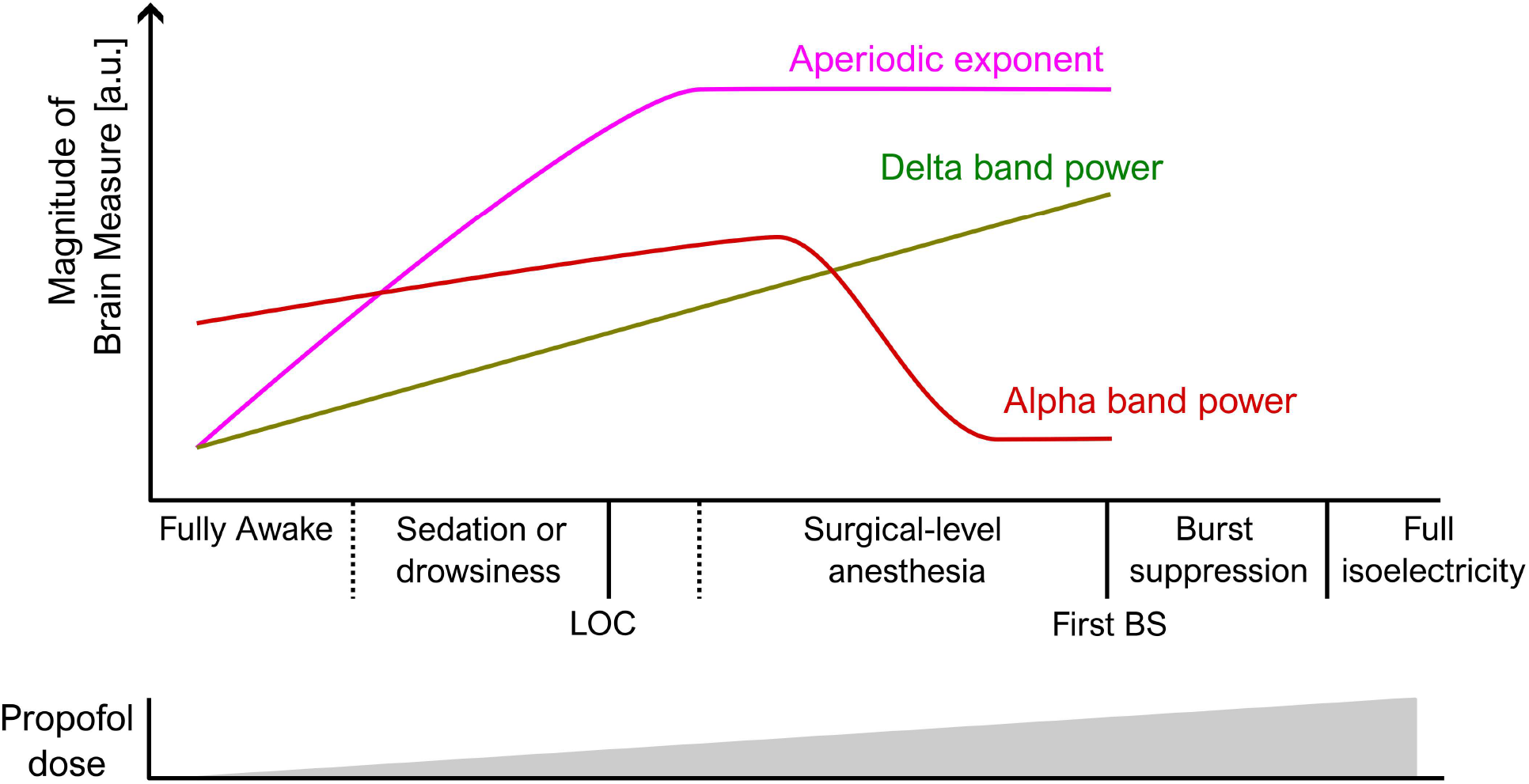
Proposed model of the relations between propofol dose and EEG measures. It shows the relation that can be observed for EEG’s aperiodic exponent, delta band power and alpha band power. Aperiodic exponent displays the most robust difference between awake and surgical-level anaesthesia, however, at surgical-level anaesthesia it ceases to be informative of different intraoperative hypnotic depths. In contrast, delta band power shows differences throughout the whole range from fully awake to the deeper surgical-level hypnotic depths. Alpha power shows a bimodal relation to hypnotic depth, with a positive relation at lower depths and a negative correlation with hypnotic depth at higher surgical-level hypnotic depths. LOC stands for the moment of loss of consciousness; First BS indicates the first burst suppression.

Several human studies have shown that inhibitory GABAergic agents like propofol or halogenated gases consistently produce steeper aperiodic slopes (higher exponents) when compared to wakefulness.^22–24, 18^ Recent studies have reported that the aperiodic exponent shows a remarkable performance in classifying subject’s EEG as awake or anesthetized, ^18^ even in patients with disorders of consciousness.^25^ This is consistent with our results where aperiodic exponent showed a greater effect size (t value) when comparing baseline to anaesthesia states than any spectral band power. However, when we probed finer changes in intraoperative propofol doses, the aperiodic exponent ceased to correlate with propofol Ce, both when employing their absolute value or the relative changes between consecutive intraoperative periods (Figures 3 and 4). Comparisons between EEG measures at different hypnotic depths are seldom conducted. A notable exception is the work by Zhang and cols. (2023) where they use a slow propofol infusion protocol to test the reliability of the aperiodic exponent as a tracker of hypnotic depth. They find a reliable and mostly linear relation between aperiodic slope and propofol dose. However, their protocol was restricted to more superficial hypnotic depths, stopping propofol infusions just after loss of consciousness. Their maximum propofol Ce ranged from 1.0 to 3.2 µg*ml^-1^ with a mean of 2.24 µg*ml^-1^ (s.d. = 0.52 µg*ml^-1^). In contrast, the propofol concentrations at which we ceased to observe a correlation between aperiodic slope and propofol dose were significantly higher than those required for loss of consciousness. They were surgery-level propofol doses and started at 2.3 µg*ml^-1^ with an average of 3.27 µg*ml^-1^ propofol Ce. In this line, we see our results expanding on previous work by illustrating both the solid presence of the relation between aperiodic exponent and propofol dose at superficial hypnotic depths but also the fact that this relation is no longer present at surgery-level hypnotic depths.

We observed a much stronger association between band powers and dose for the relative change in propofol Ce between consecutive time periods than for the absolute propofol Ce (Figure 3 vs Figure 4). We believe this is grounded in the fact that different patients have differential susceptibilities to anaesthetics. This means that, even when controlling for pharmacokinetics, a given propofol Ce can have different effects in terms of altering brain dynamics thus driving patients to different hypnotic depths.^15^ Interestingly, the level of individual susceptibility has been recently related to baseline levels of brain activity complexity ^26^ which could be an avenue to explore in the future.

In line with previous results, here we report the EEG’s aperiodic exponent as a strong correlate of changes in cortical excitation/inhibition in ranges near the point of loss of consciousness. However, this relation was not present at higher surgery-level propofol doses. Several recent works have found that this association between aperiodic exponent and cortical activity balance is not absolute. Salvatore and cols. (2024) report that only some drugs affect aperiodic exponents as expected (GABA-A agonists), but others do not (GABA-A antagonists). Similarly, in this context Ameen and cols. (2025) ^27^ argue that “the association between the [aperiodic] exponent and E/I balance is necessarily more nuanced than a one-to-one mapping”. They note that this relation depends on several parameters of model fit, and the type of data. ^28, 29^ Here we complement this by circumscribing this relation to a certain range of excitatory/inhibitory balances. Limitations of the present study include the relatively low number of patients and the fact that patients were under surgery which impeded complete control over what drugs were administered.

